# VODKA2: A fast and accurate method to detect non-standard viral genomes from large RNA-seq datasets

**DOI:** 10.1101/2023.04.25.537842

**Authors:** Emna Achouri, Sébastien A. Felt, Matthew Hackbart, Nicole S. Rivera-Espinal, Carolina B. López

**Affiliations:** Department of Molecular Microbiology and Center for Women Infectious Disease Research, Washington University School of Medicine, St Louis, MO, USA

## Abstract

During viral replication, viruses carrying an RNA genome produce non-standard viral genomes (nsVGs), including copy-back viral genomes (cbVGs) and deletion viral genomes (delVGs), that play a crucial role in regulating viral replication and pathogenesis. Because of their critical roles in determining the outcome of RNA virus infections, the study of nsVGs has flourished in recent years exposing a need for bioinformatic tools that can accurately identify them within Next-Generation Sequencing data obtained from infected samples. Here, we present our data analysis pipeline, Viral Opensource DVG Key Algorithm2 (VODKA2), that is optimized to run on a High Performance Computing (HPC) environment for fast and accurate detection of nsVGs from large data sets.

**Availability and implementation:** VODKA2 is freely available at GitHub (https://github.com/lopezlab-washu/VODKA2)

## INTRODUCTION

RNA viruses contain populations of RNA genomes that include the standard full-length viral genomes as well as truncated and/or rearranged non-standard viral genomes (nsVGs) (Gonzalez Aparicio et al., 2022). Two of the most studied nsVGs are copy-back viral genomes (cbVGs) and internal deletion viral genomes (delVGs). CbVGs are generated when during replication of the viral RNA genome the polymerase detaches from the template strand and resumes elongation downstream by copying the nascent strand generating viral genomes with complementary ends (Lazzarini et al., 1981; Perrault and Leavitt, 1978; Re et al., 1983; Vignuzzi and Lopez, 2019) (Fig. 1A). Internal delVGs have truncations either due to the viral polymerase skipping portions of the genome or the viral genome recombining to eliminate portions of the genome (Davis et al., 1980; Nomoto et al., 1979; O’Hara et al., 1984; Perrault and Semler, 1979; Vignuzzi and Lopez, 2019) (Fig. 1A). nsVGs have three well characterized functions that shape virus infection outcome: interference with standard viral genome replication, immunostimulation, and establishment of viral persistence (Vignuzzi and Lopez, 2019). Despite their biological significance, the study of nsVGs has been limited by their vast diversity and the limited presence of unique sequences that differentiate nsVGs from the standard viral genome. Although several bioinformatic tools have been created for the identification of nsVGs from NGS data sets (Beauclair et al., 2018; Bosma et al., 2019; Boussier et al., 2020; Jaworski and Routh, 2017; Olmo-Uceda et al., 2022; Routh and Johnson, 2014; Timm et al., 2014), we show in this work that these tools are not optimized for the identification of cbVGs and often lead to mistakes in the identity of the reported cbVG species. Furthermore, these tools are not designed to efficiently handle large data sets, leading to lengthy processing times. Therefore, there is a need for bioinformatics approaches that can accurately and efficiently detect and characterize cbVGs from large NGS data sets. Our group previously developed and utilized an algorithm specifically focused on the detection of cbVGs (Felt et al., 2022; Felt et al., 2021; Sun et al., 2019). Here we present a substantially updated version of this tool, the Viral Opensource DVG Key Algorithm 2 (VODKA2), that is optimized to run on a High Performance Computing (HPC) environment for fast and accurate detection of nsVGs.

**Figure 1.**
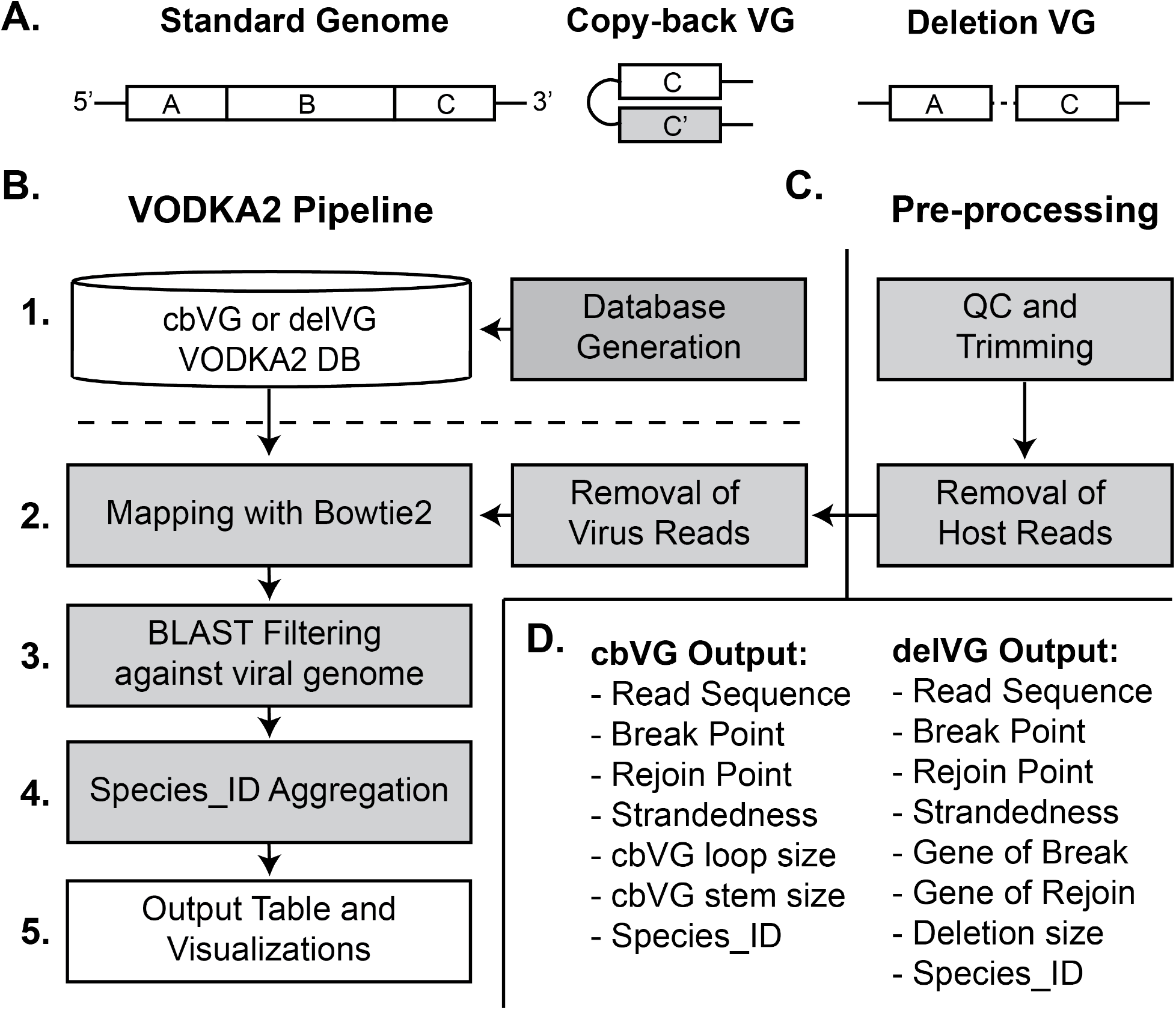
Description of the VODKA2 process. (A) Schematic of a standard viral genome (VG) containing 3 genes: A, B, and C, a copy-back VG containing gene C and the complement C’ end, and a deletion VG that has the B gene removed. (B-D) Brief descriptions of the steps and output files of the VODKA2 bioinformatics pathway. Gray boxes indicate bioinformatics programs/processes and white boxes indicate output files. (B) The steps of the VODKA2 pipeline that are discussed in the text. (C) Pre-processing steps of RNA-sequencing reads to obtain non-Host Reads.

## RESULTS

### Pipeline Overview

VODKA2 detects cbVG junctions from specific regions of the viral genome and quantifies the number of RNA-sequencing reads that contain such cbVG junction with enhanced accuracy. The VODKA2 pipeline proceeds in the following 5 steps (Fig. 1B):

#### 1. NsVG database generation

a database of all potential nsVG junctions is created based on a reference standard viral genome selected by the user. A nsVG junction is defined as the connected nucleotides where the viral polymerase breaks from the viral genome and then subsequently rejoins at either the nascent strand (copy-back genomes) or further downstream of the template (deletion genomes) to generate a continuous break/rejoin sequence. To generate the database of all potential cbVGs or delVGs, all possible combinations of break and rejoin positions together with *N* nucleotides upstream of break and downstream of rejoin are first extracted from the reference genome (*N* should be at least equal to the sequencing read size of the dataset to be analyzed). The sequence downstream of rejoin is maintained for delVG junction sequences (region A of theoretical delVG shown in Fig. 1A) or reverse complemented for cbVG junction sequences (region C’ of theoretical cbVG in Fig. 1A) before being concatenated with the sequence upstream of break (region C in Fig. 1A). Further description of the generation of the nsVG databases are detailed in the Suppl. Fig. 1. The theoretical nsVG junction sequences have a length of 2\**N*, allowing for full mapping of potential nsVG junction reads during step 2 described next.

#### 2. Mapping reads to nsVG database with Bowtie2

Before running samples through VODKA2, we recommend trimming sequencing adapters and removing host reads from the raw sequencing data (see Fig. 1C). Reads corresponding to any known co-infecting virus or contaminant organism can also be removed to reduce the size of input data, hence increasing the speed of analysis. The VODKA2 process initiates by eliminating any viral read that aligns perfectly with the standard viral genome (using Bowtie2 with default parameters). These preliminary filters help reducing the data volume and minimizing false positives, as some host and viral sequences could be mistakenly identified as non-standard viral genome (nsVG) junctions. Mapping against the VODKA2 nsVG database, which is the key step of our method, is performed with Bowtie2 set to lenient settings (option -mp 0,0) in order to minimize mismatch penalties due to viral mutations (e.g. nucleotide substitutions, insertions and/or deletions). We designed the algorithm to report any read that would match exactly 30 nucleotides around the junction (15 on each side), but any mismatch outside of that region is allowed in order to maximize the chances of detecting all nsVG junctions reads and avoid false negatives. This approach assures high nsVG detection sensitivity and generates a comprehensive preliminary list of possible reads mapped to nsVG junctions for each dataset.

#### 3. Filtering possible junction reads with BLAST

To ensure that no false positive reads are part of the output, VODKA2 runs an additional BLAST alignment of all possible nsVG reads from step 2 against the viral reference genome. Any potential nsVG read that does not meet the following criteria is discarded: (i) the read must match exactly 2 separate regions of the reference viral genome (reported as “alignment ranges” in BLAST output). Sequences mapped to the VODKA2 database during step 2 but showing ambiguous BLAST alignment (more than 2 ranges) are removed. (ii) The break/rejoin positions shown by the 2 BLAST ranges must be consistent with step 2 output (i.e. within 5 nucleotides). (iii) The last requirement is nsVG subtype specific. For delVGs, the 2 ranges must map to the same strand of the viral genome. For cbVGs, the 2 ranges must match opposite strands of the genome. Upon fulfillment of the three requirements, the read is designated as a true nsVG junction read.

#### 4. NsVG species aggregation

Among the confirmed nsVG reads, some nsVG junctions share similar sequences, but the reported break/rejoin junction is shifted to a nearby position. This shift often occurs if the break/rejoin junction contains identical sequences flanking the break and rejoin positions. NsVG reads can be mapped to any position within these sequences. This phenomenon was already reported by others (Beauclair et al., 2018; Routh and Johnson, 2014) and can potentially bias quantification of nsVGs junction reads. To provide more accurate nsVG read counts at these shifted junctions, similar reads are grouped into nsVG “species” based on two criteria: (i) the predicted nsVG size that is calculated based on the junction break and rejoin positions in the reference genome must be identical for all nsVG reads within the same group, (ii) break position shifts must be of 5 nucleotides or less. A new “Species_ID” is then generated to represent each group of similar nsVG reads. The Species_ID is called as the most represented break (and its corresponding rejoin) in the group (Suppl. Fig. 2). While this aggregation process provides a more accurate representation and quantification of nsVG junctions present in a sample, the aggregation settings can be altered depending on the question asked.

#### 5. Output Table and Visualizations

VODKA2 provides abundant output data for the detected nsVG species (Fig. 1D). For both cbVGs and delVGs, the output file includes the read ID, the sequence of the read, the break point, the rejoin point, the strandedness of the read (SAM flag), the predicted size of the nsVG and the aggregated Species_ID. For cbVGs only, the output file also includes the theoretical length of the cbVG loops and stems. For delVGs only, the output file includes the deletion size, the viral gene where the break occurs, and the viral gene where the rejoin occurs. Using this output, multiple types of analyses can be performed to study the abundance and diversity of cbVGs and delVGs.

### VODKA2 detects simulated cbVGs with high accuracy

In order to assess the accuracy of VODKA2 in detecting cbVGs, we performed a comparative analysis against state-of-the-art methodologies. We created 10 artificial cbVG sequences *in silico*, based on the respiratory syncytial virus (RSV) G gene (KC731482.1), these are detailed in Table 1. This gene contains a 72 nt repetition within its sequence, posing an additional challenge for the detection of nsVGs as it is expected to impact sequence alignments. The simulated sample was aligned against the RSV G gene reference sequence (KC731482.1) and the non-viral reads were then analyzed with VODKA2, ViReMa (Sotcheff et al., 2023)and DI-tector (Fig. 2A). VODKA2 and DI-tector detected all of the artificial cbVG junction regions without generating any false positive (Table 1 and Fig. 2A). ViReMa also detected all of the artificial cbVGs but in addition reported a false positive species. However, the Break/Rejoin positions of the cbVG junctions reported by both ViReMa and DI-tector (respectively referred as Donor/Acceptor sites and Breakpoint/Reinitiation sites) were incorrect and can mislead the user. As shown in Figure 2A, the Break and Rejoin positions of the cbVG species seem to be systematically inverted by both tools. In the case of ViReMA, this can be explained by the fact that the software was originally designed to detect recombinations between donor and acceptor genomes of the same orientation. Hence, any recombination event between a positive and a negative strand and viceversa is called a copy-back, without clearly identifying which of the donor or acceptor site is the break (or rejoin) position of a cbVG(Sotcheff et al., 2023). Regarding DI-tector, although it also seems to be biased to the opposite sense, 8 cbVG species out of 10 are reported in both orientations. This phenomenon has already been observed and reported by the authors of DI-tector and suspected to be false positives (Beauclair et al., 2018). The VODKA2 approach presents the advantage of avoiding such ambiguous reporting as the pre-built database will constrain the putative cbVG to a specific orientation. In other terms, the Break and Rejoin reported by VODKA2 are controlled by the cbVG reference sequences and cannot be distort by a misinterpretation of the sequencing reads orientation.

**Table 1.** Simulation of RSV G copy-back and deletion sequences.

| Type of NsVG | Break Position | Rejoin Position | Genome Coverage | Total Reads | NsVG Junction Reads | VODKA2 | ViReMa | DI-tector |
| --- | --- | --- | --- | --- | --- | --- | --- | --- |
| CB | 150 | 795 | 330 | 1081 | 234 | 219 | 227 | 212 |
| CB | 261 | 404 | 180 | 757 | 44 | 30 | 44 | 44 |
| CB | 274 | 652 | 70 | 234 | 31 | 31 | 31 | 17 |
| CB | 299 | 799 | 40 | 111 | 31 | 31 | 30 | 30 |
| CB | 391 | 669 | 60 | 174 | 35 | 24 | 35 | 35 |
| CB | 590 | 810 | 50 | 98 | 20 | 18 | 20 | 20 |
| CB | 594 | 701 | 30 | 63 | 17 | 16 | 17 | 17 |
| CB | 689 | 849 | 140 | 184 | 74 | 74 | 74 | 74 |
| CB | 822 | 915 | 20 | 13 | 6 | 6 | 6 | 6 |
| CB | 832 | 930 | 80 | 46 | 21 | 21 | 21 | 21 |
| **CB** | **300** | **869** | - | - | - | - | **1** | - |
| DEL | 190 | 472 | 160 | 363 | 71 | 71 | 71 | 71 |
| DEL | 212 | 800 | 350 | 439 | 135 | 135 | 131 | 128 |
| DEL | 249 | 549 | 170 | 376 | 53 | 53 | 52 | 53 |
| DEL | 251 | 281 | 70 | 217 | 26 | 26 | 25 | 0 |
| DEL | 273 | 562 | 490 | 1100 | 174 | 53 | 174 | 164 |
| DEL | 368 | 522 | 100 | 269 | 33 | 33 | 31 | 32 |
| DEL | 441 | 751 | 110 | 239 | 23 | 23 | 22 | 22 |
| DEL | 470 | 497 | 110 | 342 | 1 | 0 | 0 | 0 |
| DEL | 535 | 672 | 180 | 495 | 25 | 25 | 25 | 25 |
| DEL | 692 | 747 | 230 | 695 | 22 | 21 | 20 | 8 |
| **DEL** | **212** | **872** | - | - | - | - | **2** | - |

Green cells indicate results that are identical to the expected. Red cells indicate false positives.

**Figure 2.**
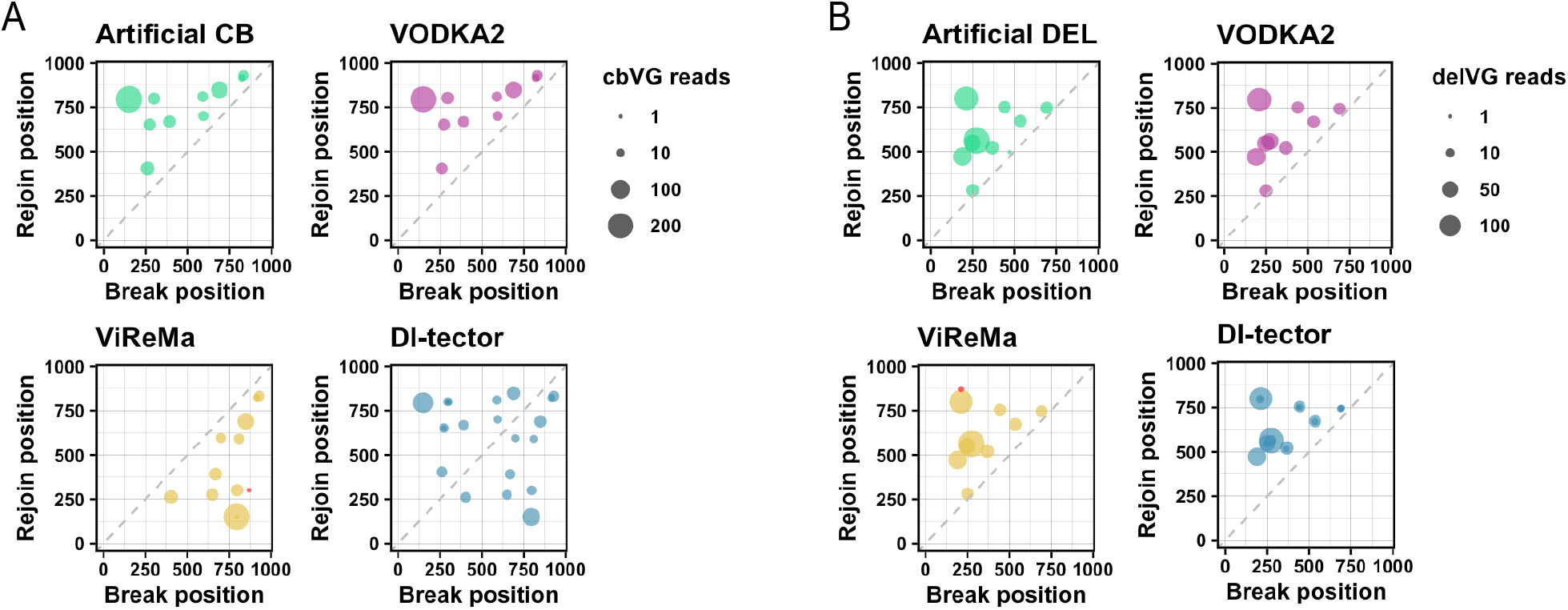
Detection of simulated cbVG and delVG reads. (A+B) Representation of (A) cbVG and (B) delVG break and rejoin points reported by VODKA2 (pink), ViReMa (yellow) and DI-tector (blue). Expected results are shown in green. Size of dots is dependent on the number of detected read at each break/rejoin junction. False positive reads are shown in red.

### VODKA2 can detect delVGs

In addition to cbVGs, it is possible to use VODKA2 to detect deletions by generating a database of all putative delVGs of the specified reference genome. To illustrate this new feature, we analyzed 10 artificial delVGs based on the RSV G gene. Similarly to the results obtained with the simulated cbVG data, all expected delVG species were correctly identified, showing again the specificity of this method. Observably, VODKA2 reported slightly fewer reads than DI-tector and ViReMA for some particular cbVG and delVG species (Fig 2B and Table 1) and it is unclear at the moment what is the couse of this.

### VODKA2 detects cbVGs from NGS data with high accuracy and efficiency

With the goal of demonstrating the cbVGs detection efficiency of VODKA2 in real RNA-seq data, we analyzed data from a sample of cells infected *in vitro* with Sendai virus (SeV), which is a paramyxovirus of 15,384 nucleotides that is known to produce a single major cbVG species of length 546 nucleotides (Sun et al., 2019), with the Break and Rejoin occurring around positions 14,930 and 15,290 of the reference genome. VODKA2 expeditiously processed the sample and reported 7,138 validated cbVG junction reads, all having a Break and Rejoin around the expected positions for cbVG 546. ViReMa and DI-tector results also confirmed the presence of this major species but both algorithms also reported at least 1 false positive junction and required significantly increased running time compared to VODKA2, which processed the sequencing sample almost 10 times faster. This comparison demonstrates the advantage of VODKA2 to fast and reliably detect cbVGs in large datasets (Fig. 3).

**Figure 3.**
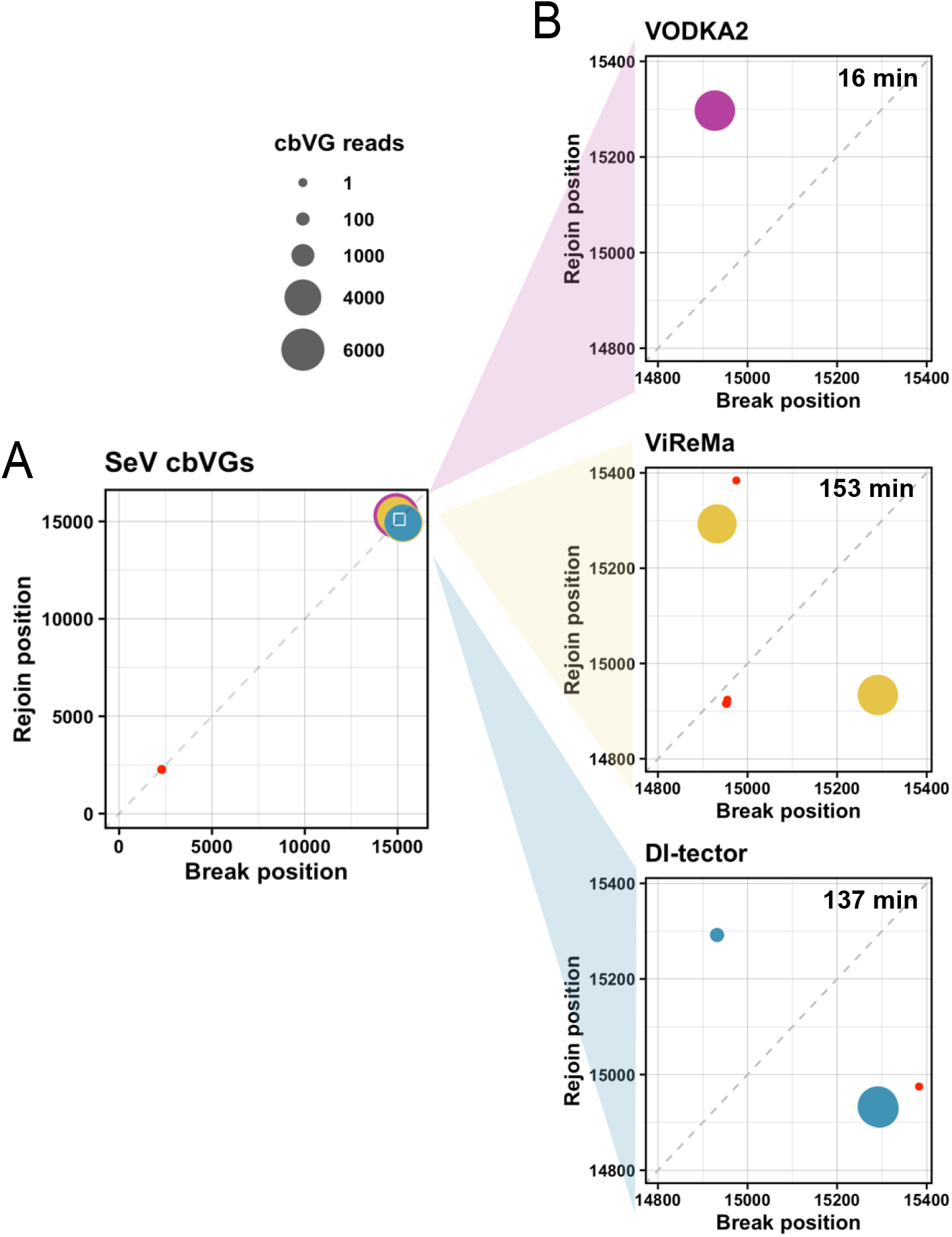
Detection of cbVG reads in SeV. Representation of cbVG break and rejoin points reported by VODKA2 (pink), ViReMa (yellow) and DI-tector (blue). Size of each point is dependent on the number of detected read at each break/rejoin species. (A) Shows an overlay of all cbVGs detected across the whole genome by the three softwares. Red dot indicates a false positive from DI-tector. (B) Zooms of species detected within positions 14,800 and 15,384 of the reference genome for each software. False positive reads are shown in red. Processing time of each software is shown in minutes.

## DISCUSSION AND CONCLUSION

It has become clear over the past decade that nsVGs play a crucial role in virus pathogenesis (Felt et al., 2021; Penn et al., 2022; Vasilijevic et al., 2017; Vignuzzi and Lopez, 2019; Zhou et al., 2022). As a consequence, there has been a big push in the nsVG field to develop NGS/bioinformatics approaches that are more sensitive, less biased, and more accurate, such as DI-tector (Beauclair et al., 2018), DVG-Profiler (Bosma et al., 2019), DG-Seq (Boussier et al., 2020), ViReMa (Jaworski and Routh, 2017; Routh and Johnson, 2014; Sotcheff et al., 2023) and DVGfinder (Olmo-Uceda et al., 2022). Our own group has previously developed a pipeline, VODKA, which is the first tool focusing on the detection of cbVGs within NGS (Sun et al., 2019). We have updated and improved this pipeline, now called VODKA2, to detect not only cbVGs but also delVGs in a time effective manner.

The development of NGS technologies offers the capacity of deeper sequencing and increasing amounts of samples within a single experiment, which raises the need for improving the running speed of bioinformatics tools for more efficient analysis of large data sets. We achieved that goal in VODKA2 and considerably accelerated the analysis process of each sample by separating the nsVG database generation step. A database is generated only once for each reference genome and remains available for any further sample analysis, saving precious processing time when running larger datasets. Moreover, contrary to other common methods such as ViReMa and DI-tector, VODKA2 database offers the benefit of calling nsVG junctions after running a single alignment step, without having to split unmapped sequences and re-align them multiple times. Furthermore, we designed VODKA2 to easily run several samples in parallel by using a job scheduler when available, and take advantage of high performance computing resources, offering the possibility to run effectively large-scale project.

Another strong advantage of VODKA2 nsVG database approach is the control of cbVG orientation by unambiguously identifying the Break and Rejoin positions, whereas both ViReMa and DI-tector report misleading information that can be confusing for the user, unless running further analysis of each reported junction read to identify the correct sense. Yet, as cbVGs have been shown to be produced only during replication of negative-sense RNA viruses, VODKA2 focuses by default on the cbVGs generated from the 3’ ends of the provided reference genome. If looking to detect cbVGs generated from the opposite end, e.g. when working with a positive-sense RNA virus, we suggest using the reverse sequence of the reference genome to generate the cbVG database and running the analysis.

VODKA2 can analyze large NGS datasets and generates accurate and detailed outputs for further analysis of cbVGs and delVGs. We show that the new filtering process reduces the risk of detecting false positives, increasing the specificity of VODKA2. Altough we observed a slight decrease of VODKA2 sentivity as compared to ViReMa and DI-tector, that is at the benefit of a 10 time fold increase of speed. Taken together, VODKA2 is a valuable tool for the fast detection of nsVG species providing data that set the stage for understanding better the biology of nsVG amongst different RNA virus populations in large-scale studies.

## MATERIAL AND METHODS

### Simulated nsVG sequencing data

We used RSV reference genome (KC731482.1) and generated random numbers from a range of 15 to 951 to create artificial copy-back and deletion sequences from gene G. Novaseq150 bp paired end reads were simulated using InSilicoSeq v1.5.3 with default parameters and random coverage values (Gourle et al., 2019).

### RNA-Seq data pre-processing

Raw sequencing reads were trimmed of Illumina adaptors using Cutadapt (v2.5) and analyzed by FastQC (v0.11) to ensure quality of sequencing reads. The reads mapping the human genome (GRCh38) were removed from the datasets using Bowtie2 (v2.4.1).

### nsVG detection with VODKA2

VODKA2 nsVG databases were created using VODKA2 *genome_to_new_fasta* step with read size set to 150bp. We used the whole viral reference genome size for all databases: 966nt from RSV gene G reference sequence were used to generate the cbVG and delVG databases. With Sendai virus being much larger, we used 400GB RAM memory to generate cbVG and delVG databases from the whole reference genome (15,384). The output of this step is a multi FASTA file containing all theoretical junction region sequences, which was then indexed for Bowtie2 (using Bowtie2-build v2.4.1).

VODKA2 analysis was performed according to the recommendations on GitHub. VODKA2, ViReMa and DI-tector analyses were run using either 10GB for the simulated nsVG data or 150GB for the SeV infected sample. All scatterplots shown in this work were generated using R v4.3.0.

### Cell line and virus

A549 cells (human type II pneumocytes cells, ATCC CCL-185) were cultured at 37°C with 5% CO2 with Dubelcco’s modified Eagle’s medium (DMEM; Invitrogen) supplemented with 10% fetal bovine serum (FBS), 50 ng/ml gentamicin (Thermofisher), 2 mM L-glutamine (Invitrogen), and 1 mM sodium pyruvate (Invitrogen). A549 cells were treated with mycoplasma removal agent (MP Biomedicals, 093050044) and routinely tested for mycoplasma before use. Sendai Cantell (SeV) strain was grown in 10-day-old, embryonated chicken eggs (Charles River) for 40 h as previously described (Yount et al., 2006).

### Virus infection

A549 cells were washed twice with PBS. Subsequently, the cells were exposed to SeV virus in an infectious medium, with a multiplicity of infection (MOI) of 1.5. The infectious medium consisted of DMEM supplemented with 35% bovine serum albumin (BSA; Sigma-Aldrich), penicillin-streptomycin (Invitrogen), and 5% NaHCO3 (SigmaAldrich). The virus and cells were incubated together at 37°C for 1.5 hours. Following the incubation period, additional infectious medium was provided to the cells. As a control, mock conditions underwent the same media change procedure but without the addition of the virus. Infected and mock cells were incubated at 37°C and 5% CO2 for 16 hours. Supernatant were collected of cells and RNA was extracted by TRIzol/chloroform extraction (Thermofisher, 15596018).

### RNA-Sequencing

Total RNA was extracted from mock and infected cells using TriZol reagent (Life Technologies). The concentration of the extracted RNA was measured using a Thermo Scientific™ NanoDrop™ spectrophotometer. RNA quality was assessed using an Agilent TapeStation or Bioanalyzer (Agilent Technologies) prior to cDNA library preparation. All samples were prepared using the Illumina TruSeq Stranded Total RNA Library Prep Kit with Ribo-Zero Gold. SeV samples were run on an Illumina NextSeq 500 to generate 150 bp, paired-end reads, resulting in ∼ 41 million reads per sample. SeV samples were run on an NextSeq 550 to generate 150 bp, pairerd-end reads, resulting in ∼ 64 to 91 million reads per samples. Average phred quality score of samples were ∼ 34.

## SUPPLEMENTAL MATERIAL

Supplemental material is available for this article.

## ACKNOWLEDGEMENTS

We want to thank Gregory R Grant and Eun Ji Kim for initial VODKA.

## FUNDING INFORMATION

This works was supported by the US National Institutes of Health National Institute of Allergy and Infectious Diseases AI137062 and AI134862 to CLB, NSF-GRFP DGE2139839 to NSR, and T32-007317 to MH.

## Notes

https://github.com/lopezlab-washu/VODKA2

### Competing Interest Statement

The authors have declared no competing interest.

### Summary of Updates

We now include a comparison of VODKA2 with other state of the art similar tools

