## supplemental tables for "VODKA2: A fast and accurate method to detect non-standard viral genomes from large RNA-seq datasets"

**Supplementary Information**

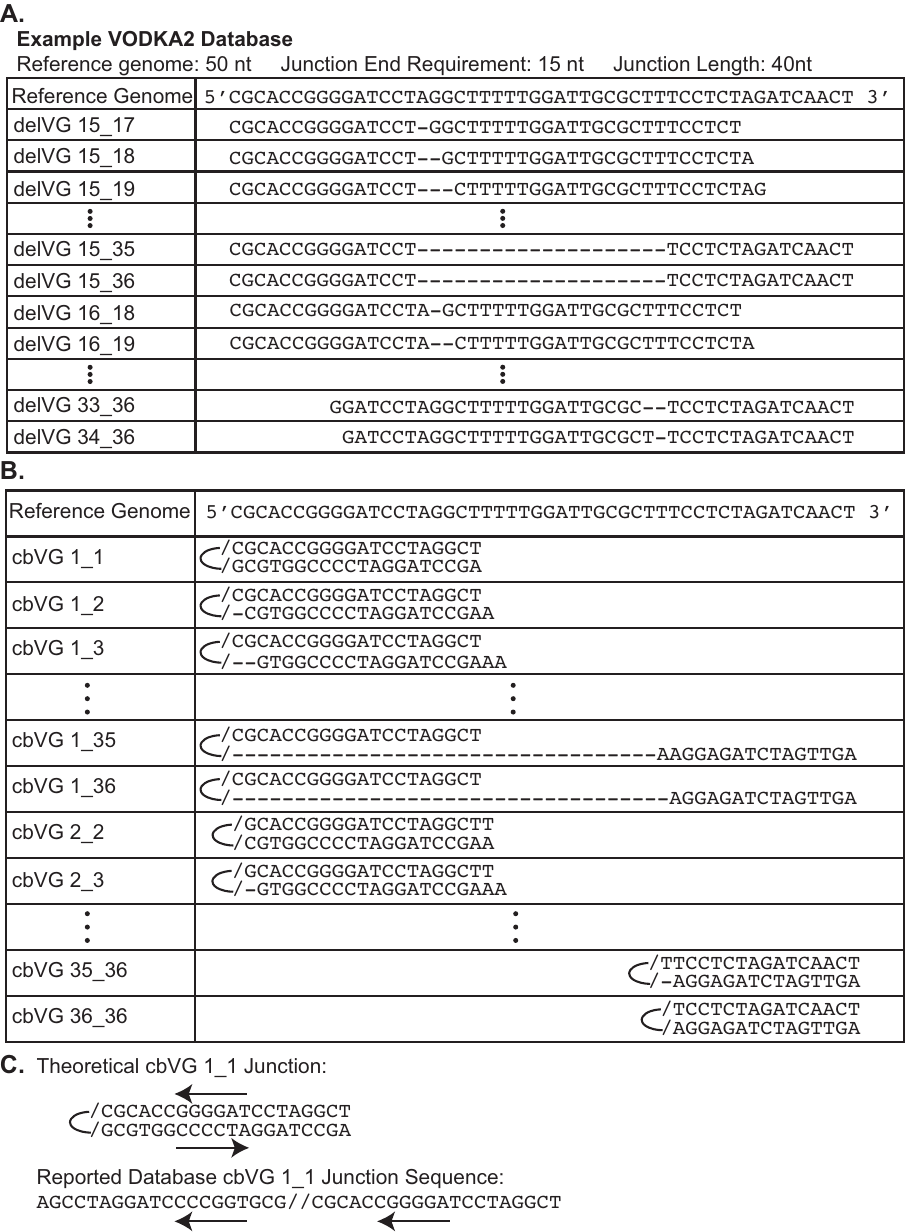

**Supplemental Figure 1: Example of VODKA Database Generation.** A) Schematic for VODKA2 delVG database generation. B) Schematic for VODKA2 cbVG database generation. C) Orientation and readout of cbVG junction in VODKA2 database.

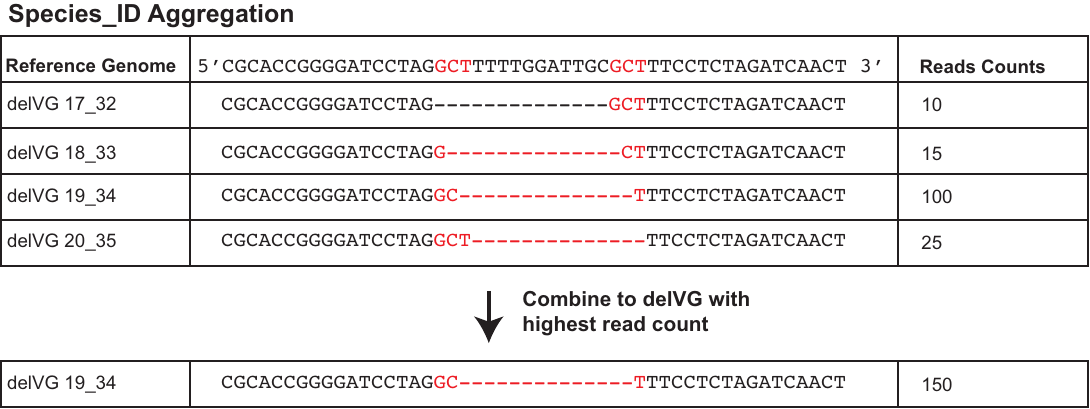

**Supplemental Figure 2: Example of Species_ID Aggregation.** Species_ID pooling for delVG 17_32, delVG 18_33, delVG 19_34, and delVG 20_35, where the delVG read counts are summed and assigned to the mode species within the group.

**Supplemental Table 1: List of ns VG databases used in this work**

|  | **reference sequence length** | **read length** | **VODKA2 db** | | | |
| --- | --- | --- | --- | --- | --- | --- |
|  |  |  | **type** | **cbVG junctions** | **# bases** | **file size** |
| **RSV G gene** | 966 | 150 | CB | 453,628 | 127,793,488 | 130M |
| **IAV PB1** | 2341 | 250 | DEL | 2,671,516 | 1,214,587,336 | 1.2G |
| **IAV PB2** | 2341 | 250 | DEL | 2,671,516 | 1,214,587,336 | 1.2G |
| **IAV M** | 1027 | 250 | DEL | 497,503 | 198,281,263 | 198M |
| **IAV NS** | 890 | 250 | DEL | 370,230 | 142,115,510 | 142M |

**Supplemental Table 2: List of artificial cbVGs from RSV gene G and VODKA2 analysis results.**

| **cbVG junction** | | **3. Length** | **4. Coverage** | **raw reads** | | **results** | |
| --- | --- | --- | --- | --- | --- | --- | --- |
| **1. Break** | **2. Rejoin** |  |  | **5. Expected** | **6. Simulated** | **7. Standard** | **8. Junction** |
| 150 | 795 | 989 | 330 | 2175 | 2162 | 1915 | 236 |
| 261 | 404 | 1269 | 180 | 1522 | 1514 | 1439 | 60 |
| 274 | 652 | 1008 | 70 | 470 | 468 | 430 | 38 |
| 299 | 799 | 836 | 40 | 222 | 222 | 193 | 26 |
| 391 | 669 | 874 | 60 | 349 | 348 | 315 | 22 |
| 590 | 810 | 534 | 50 | 178 | 176 | 139 | 36 |
| 594 | 701 | 639 | 30 | 127 | 126 | 112 | 14 |
| 689 | 849 | 396 | 140 | 369 | 368 | 252 | 114 |
| 822 | 915 | 197 | 20 | 26 | 26 | 10 | 16 |
| 832 | 930 | 172 | 80 | 91 | 92 | 48 | 44 |

Columns 1 and 2: cbVG junctions represented by their break and rejoin positions. Column 2: cbVG length calculated as (break position – gene size) + (rejoin position – gene size) + 2, where the size of RSVG G gene is 966nt. Column 3: coverage value used for InSilicoSeq reads simulation. Column 4: number of expected reads as coverage * gene length / read length. Column 5: number of reads simulated with InSilicoSeq. Column 7: number of reads matching the standard RSV G gene sequence. Columns 8: number of reads detected by VODKA2 as cbVG junction.
